# Global brain activity links subcortical degeneration to cortical tau progressively across Braak regions over early Alzheimer’s disease stages

**DOI:** 10.64898/2026.01.23.701360

**Authors:** Yutong Mao, Baizhou Pan, Xiao Liu, the Alzheimer’s Disease Neuroimaging Initiative

## Abstract

Alzheimer’s disease (AD) is characterized by early tau pathology in subcortical neuromodulatory nuclei, followed by progressive cortical tau accumulation; however, the mechanisms linking subcortical dysfunction to cortical tau pathology remain unclear. Using multimodal neuroimaging data from the ADNI cohort, we examined how infra-slow (< 0.1 Hz) global brain (i.e., gBOLD) activity is related to the volume of the nucleus basalis of Meynert (NbM) and cortical tau accumulations in the early stages of AD. NbM degeneration was associated with reduced gBOLD activity and spatially co-localized tau accumulation, appearing in early Braak regions during the preclinical stage, i.e., cognitively unimpaired participants with abnormal CSF markers, and extending to more advanced Braak areas during the prodromal stage, i.e., mild cognitive impairment (MCI) subjects. Our findings suggest that infra-slow gBOLD activity serves as a functional neural mediator linking subcortical degeneration to cortical tau pathology, highlighting a potential functional pathway linking subcortical and cortical pathology in early AD.

## Introduction

Alzheimer’s disease (AD) is a progressive neurodegenerative disorder characterized by the accumulation of amyloid-β (Aβ) plaques and hyperphosphorylated tau tangles. According to the amyloid cascade hypothesis, Aβ accumulation initiates a pathological cascade that includes tau aggregation, neurodegeneration, and cognitive decline^1–3^. Growing evidence suggests that tau pathology is more closely associated with neuronal loss and clinical symptoms^4–6^. Cortical tau aggregates into neurofibrillary tangles following a specific spatial trajectory, beginning in the entorhinal cortex and progressing toward widespread neocortical areas, as classically described by the Braak staging framework^7,8^.

The cholinergic hypothesis, another classic theory of AD etiology, proposes that degeneration of basal forebrain cholinergic neurons plays a central role in AD pathophysiology. The nucleus basalis of Meynert (NbM), the principal source of cortical cholinergic innervation, is especially vulnerable in AD and undergoes early and progressive degeneration^9–12^. The association between the cholinergic dysfunction and Aβ accumulation in AD has long been recognized^13–17^, however, the directionality of this relationship had been a focus of debate^14,15^. Beyond Aβ, emerging evidence indicates that tau pathology is also closely linked to the cholinergic innervations^18^. Histological data consistently show tau pathology in NbM cholinergic neurons at the earliest stages of AD^19–22^, and neuroimaging studies have associated smaller NbM volumes with cortical tau burdens^23^. Importantly, the NbM atrophy preceded and predicted the degeneration of the entorhinal cortex^24,25^. These findings suggest early NbM change and its association with later cortical pathology in AD, yet the mechanisms underlying this relationship remain elusive.

Recent research on resting-state, infra-slow (<0.1 Hz) global brain activity positioned it as a potential functional mediator linking cholinergic dysfunction to cortical pathologies in AD. This highly structured global brain activity manifests as global waves propagating across cortical hierarchies in human fMRI and monkey electrocorticography (ECoG)^26–28^, whereas as spiking cascades of sequential activations in mouse neuronal recordings^29^. Neuromodulatory systems, particularly the cholinergic system, are critically involved in this global brain dynamic, as evident by strong and specific de-activation in their key subcortical nodes, such as the NbM^26,30^, at peaks of global mean blood-oxygenation-level-dependent (gBOLD) signal, i.e., the fMRI measure of global bran activity. Importantly, pharmaceutically de-activating one-side NbM in monkeys suppressed the gBOLD activity of the ipsilateral side^31^. Meanwhile, reduced gBOLD activity has been associated with cortical Aβ/tau accumulations in AD^32–34^, presumably because of its role in perivascular clearance via vasomotion and cerebrospinal fluid (CSF) dynamics regulated by subcortical neuromodulatory nuclei^35,36^. Thus, gBOLD activity may act as a functional mediator potentiating subcortical NbM changes to cortical pathology. This hypothesis is supported by a recent multi-modal study demonstrating that gBOLD activity is associated with cortical cholinergic activity in humans via PET-MRI; furthermore, lesions of NbM cholinergic neurons in mice reduced gBOLD activity and impaired cortical waste clearance^37^. Despite these findings, the three-way relationship among NbM degeneration, gBOLD activity, and cortical tau spreading over early AD progression remains to be elucidated.

Here we leverage multi-modal neuroimaging data from the Alzheimer’s Disease Neuroimage Initiative (ADNI) to investigate changes in, and relationships among, NbM volume, gBOLD activity, and tau accumulations during the preclinical and prodromal stages of AD. We found that smaller NbM volume was associated with weaker gBOLD activity and greater cortical tau accumulation, which co-localized spatially. These associations emerged in early Braak regions in cognitively unimpaired individuals with abnormal CSF Aβ/tau markers and progressed to advanced Braak areas in those with mild cognitive impairment (MCI). Longitudinal analyses further suggest a directional pattern whereby NbM-related reductions in gBOLD activity predicted subsequent increases in cortical tau pathology. Together, these findings suggest that infra-slow gBOLD activity may represent brain dynamics that mediate the effect of subcortical cholinergic degeneration on cortical tau accumulation during the early stages of AD.

## Results

### Demographics of subjects

We analyzed data from 75 ADNI-3 participants (73.4 ± 7.4 years, 38 females; **Table 1**) without a clinical diagnosis of AD, each of whom has resting-state fMRI and [¹□F] flortaucipir PET data from at least 2 visits with a minimal interval of 9 months. The cohort includes 37 cognitively normal (CN) individuals, 3 with significant memory concern (SMC), and 35 with mild cognitive impairment. The CN and SMC participants were combined into a cognitively unimpaired (CU) group, which was then divided stratified by CSF p-Tau/Aβ42 ratio into abnormal (≥ 0.028, *N* = 17; CU-aCSF) and normal (<0.028, *N* = 23; CU-nCSF) subgroups^38^. The CU-aCSF group is significantly older than both the CU-nCSF (*p =* 0.006) and MCI (*p =* 0.01) groups. This is of minimal concern, as all major analyses were conducted within each group and adjusted for age and sex effects, which will not be reiterated in subsequent sections.

**Table 1.**
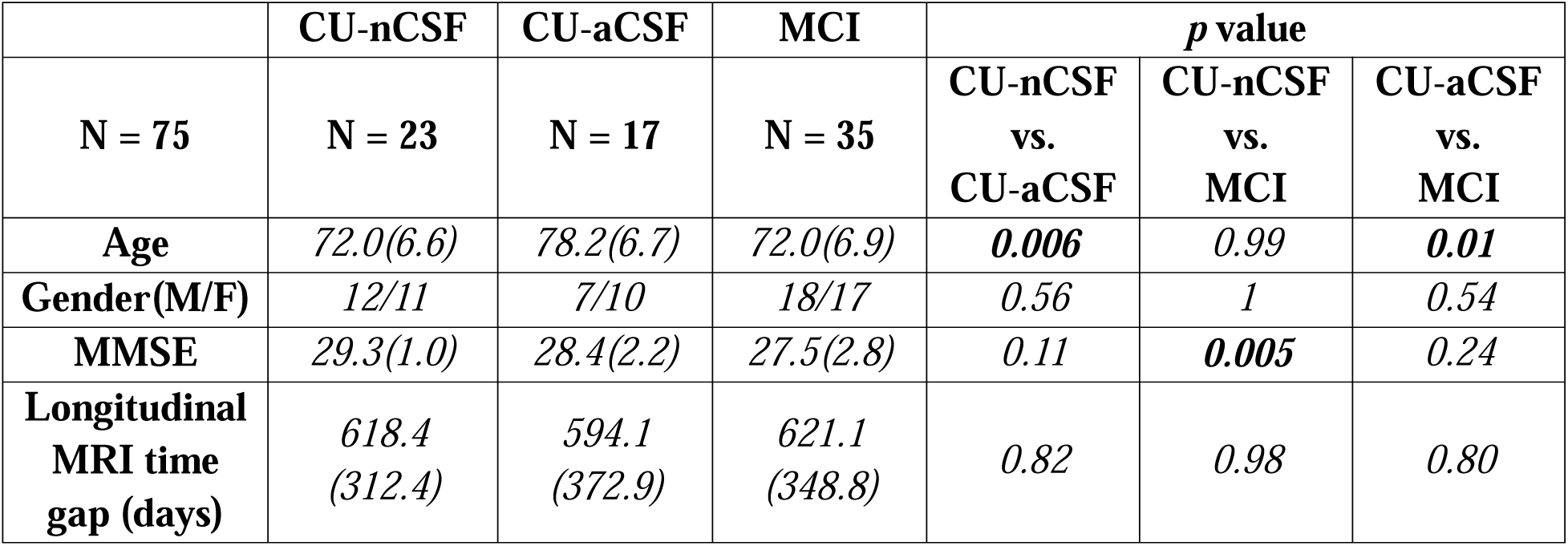
Demographic and clinical characteristics of the study participants. . Data are presented as mean (standard deviation) unless otherwise indicated. Pairwise group comparisons were performed using two-sided two-sample t-tests for continuous variables and Fisher’s exact test for categorical variables (gender). Statistically significant results are shown in **bold**. **Abbreviations:** M/F, male/female; CU-aCSF, cognitively unimpaired with abnormal CSF biomarkers; CU-nCSF, cognitively unimpaired with normal CSF biomarkers; MCI, mild cognitive impairment. MMSE, Mini-Mental State Examination.

|  | CU-nCSF | CU-aCSF | MCI | *p* value |  |  |
| --- | --- | --- | --- | --- | --- | --- |
| N = 75 | N = 23 | N = 17 | N = 35 | CU-nCSF<br>vs.<br>CU-aCSF | CU-nCSF<br>vs.<br>MCI | CU-aCSF<br>vs.<br>MCI |
| Age | 72.0(6.6) | 78.2(6.7) | 72.0(6.9) | **0.006** | 0.99 | **0.01** |
| Gender(M/F) | 12/11 | 7/10 | 18/17 | 0.56 | 1 | 0.54 |
| MMSE | 29.3(1.0) | 28.4(2.2) | 27.5(2.8) | 0.11 | **0.005** | 0.24 |
| Longitudinal<br>MRI time<br>gap (days) | 618.4<br>(312.4) | 594.1<br>(372.9) | 621.1<br>(348.8) | 0.82 | 0.98 | 0.80 |

### Associations between the NbM volume and cortical tau pathology

Smaller NbM volumes were significantly associated with higher mean cortical tau standardized uptake value ratios (SUVR) (_ρ_ = −0.49, *p* = 1.16×10^-5^) across all 75 participants; however, no such association was observed for a control region, the primary sensory cortex (PSC, area 3a) (_ρ_ = −0.11, *p* = 0.37) (**Fig.1A**). The negative NbM-tau association remains significant in both CU-aCSF (_ρ_ = −0.52, *p* = 0.04) and MCI (_ρ_ = −0.54, *p* = 0.001) groups, but not in the CU-nCSF group (_ρ_ = −0.13, *p* = 0.55) (**Fig. 1B**). Using predictive modeling on longitudinal data^24^, we found that baseline NbM volume predicted longitudinal tau changes at follow-up in both the CU-aCSF (_ρ_ = −0.53, *p* = 0.03) and MCI groups (_ρ_ = −0.42, *p* = 0.01), though in distinct regions (**Fig. S1**), but not in the CU-nCSF group (_ρ_ = 0.01, *p* = 0.96) (**Fig. 1C**). Conversely, baseline cortical tau SUVR showed a marginal association with longitudinal NbM volume changes only in the CU-aCSF group (_ρ_ = −0.46, *p* = 0.06) but not in the MCI (_ρ_ = −0.14, *p* = 0.43) group (**Fig. 1D**). These findings support that the NbM degeneration precedes and predicts cortical tau accumulation in participants with MCI, a bi-directional relationship between the two in the CU-aCSF individuals, but the absence of any temporal relationships in the CU-nCSF group (**Fig. 1E**).

**Figure 1.**
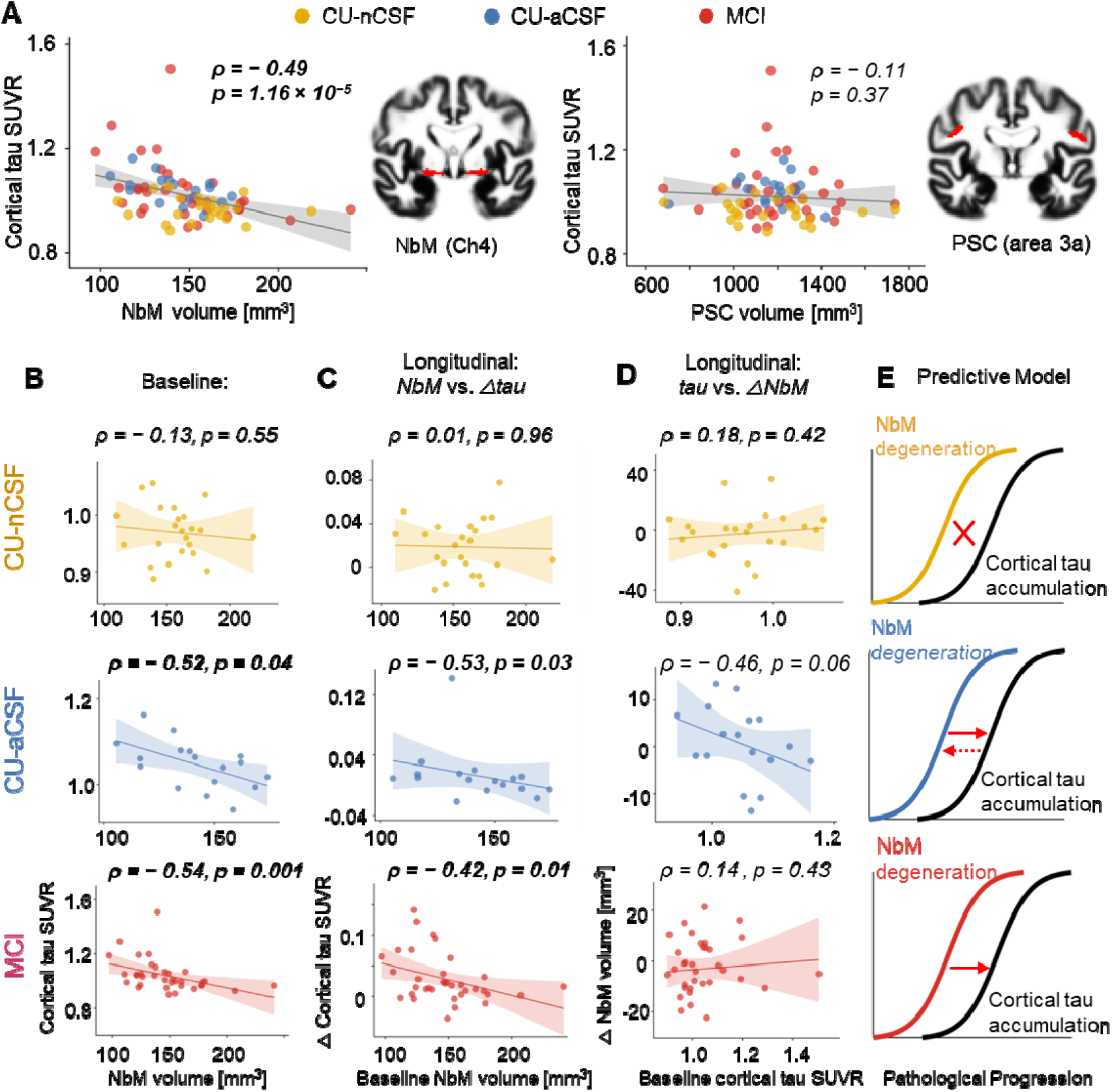
Associations between NbM volume and cortical tau burden across early AD stages. *(A)* Across all 75 participants, smaller NbM volume was significantly associated with greater cortical tau SUVR (Spearman ρ = −0.49, p = 1.16 × 10□□). In contrast, no significant association was observed between the primary sensory cortex (PSC, area 3a) volume and cortical tau (ρ = −0.11, p = 0.37). Brain overlays (red) illustrate NbM and PSC ROIs. (**B**) Group-specific analyses revealed a stage-dependent NbM–tau relationship: no association in CU-nCSF (ρ = −0.13, p = 0.55, yellow), but significant negative correlations in CU-aCSF (ρ = −0.52, p = 0.04, blue) and MCI (ρ = −0.54, p = 0.001, red). (**C**) Longitudinal analyses showed that smaller baseline NbM volume predicted more subsequent cortical tau accumulation in CU-aCSF (ρ = −0.53, p = 0.03, blue) and MCI groups (ρ = −0.42, p = 0.01, red), but not in the CU-nCSF group (ρ = 0.01, p = 0.96, yellow). (**D**) Conversely, baseline cortical tau SUVR only shows marginal correlation with NbM change in CU-aCSF group (ρ = −0.46, p = 0.06, blue). (**E**) Schematic summary of the predictive modeling results. The colored curve (yellow, blue, red) represents NbM degeneration, and the black curve represents cortical tau accumulation.

### Associations between gBOLD activity and cortical tau accumulation

We then examined associations between gBOLD activity, quantified by co-activations at gBOLD peaks^30^ (**Fig. 2A**), and cortical tau accumulation. First, gBOLD activation and cortical tau accumulation exhibited similar spatial patterns, evidenced by a significant negative correlation (_ρ_ = −0.35, *p* = 0.004, *N* = 68 parcels) between their group-mean maps. Brain regions with lower gBOLD activation exhibited greater tau accumulation (**Fig. 2B**, upper). Notably, both quantities exhibited progressive changes across sets of brain ROIs (**Table S1**) defined by Braak stages^7,8^ (**Fig. 2B**, lower). More importantly, cross-subject, region-specific correlations between gBOLD activation and tau burden exhibited distinct, group-specific patterns. In the CU-aCSF group, negative gBOLD-tau correlations were strongest in the earliest Braak regions (Braak I: _ρ_ = −0.58, *p* = 0.02; Meta-temporal: _ρ_ = −0.57, *p* = 0.02) and gradually decreased towards ROIs of later Braak stages (**Fig. 2C**; blue). In contrast, the strongest gBOLD-tau correlations were found in more advanced Braak III–IV (_ρ_ = −0.46, *p* = 0.006) and Braak V (_ρ_ = −0.48, *p* = 0.004) regions for the MCI group (**Fig. 2C**; red). No significant gBOLD-tau associations were found in any ROI sets in the CU-nCSF group (**Fig. S2**).

**Figure 2.**
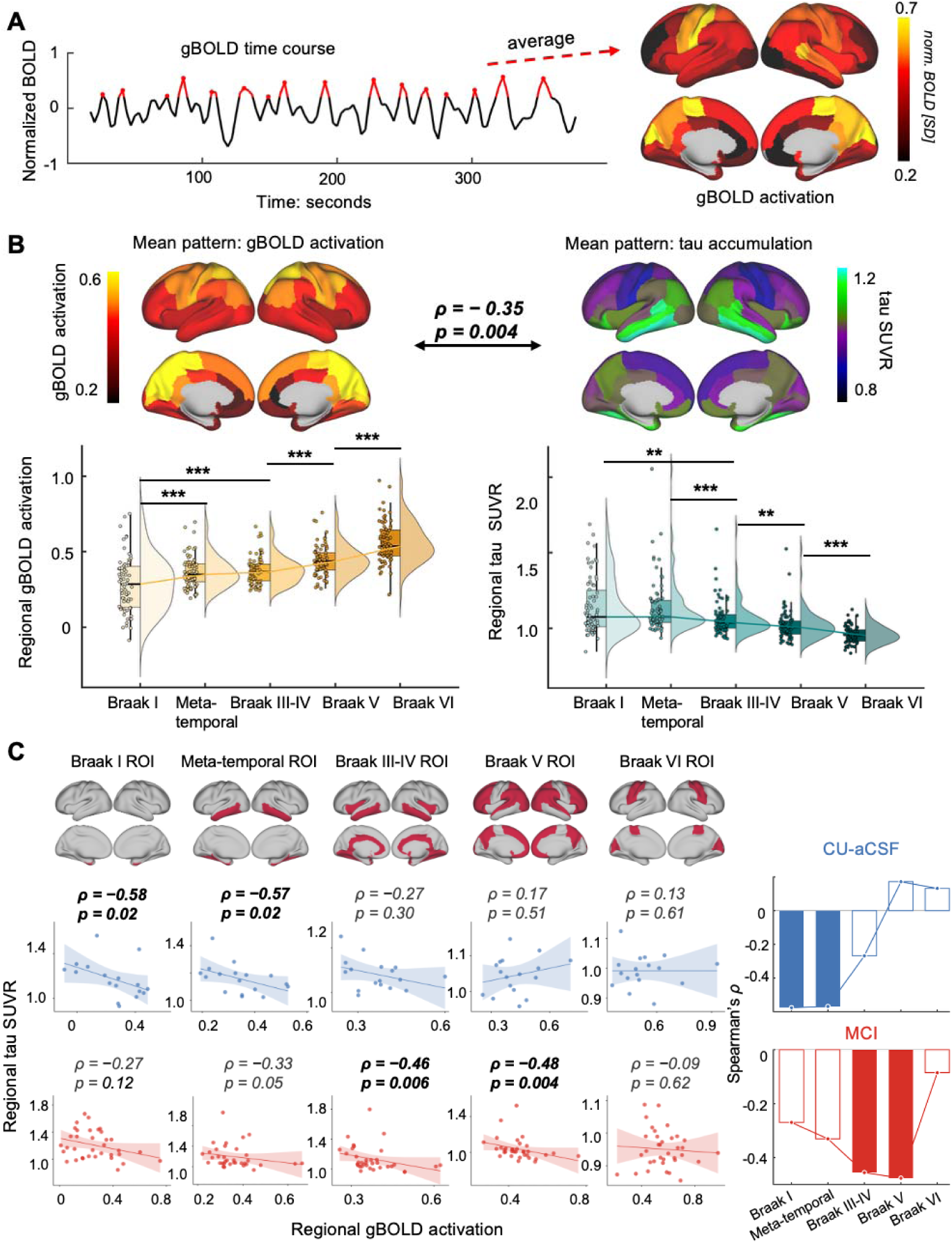
Associations between gBOLD activity and cortical tau accumulation across early AD stages. (**A**) The gBOLD activation map from a representative participant (right) was derived by averaging fMRI BOLD co-activations at gBOLD peak time points (red dots in the time series on the left)^30^. (**B**) Spatial similarity between gBOLD activation and tau accumulation. The mean spatial patterns of gBOLD activations (upper left) and cortical tau accumulation (upper right), averaged across all 75 subjects, showed a significant negative correlation (ρ = −0.35, p = 0.004). Two quantities were also summarized and compared across Braak ROI sets (lower). Asterisks indicate the statistical significance level (*: 0.01 < p < 0.05, **: 0.001 < p < 0.01, ***: p < 0.001). (**C**) Cross-subject gBOLD–tau associations across Braak ROIs in the CU-aCSF (blue, upper) and MCI (red,lower) groups. The bar plots (right) summarize correlation coefficients (ρ) across Braak regions for each group.

The correlation between NbM volume and gBOLD activation was also observed, with a similar pattern of progression from the CU-aCSF group to the MCI group (**Fig. S3**). The NbM-gBOLD correlation reached its maximum at the Braak I (_ρ_ = 0.45, *p* = 0.07) and meta-temporal ROI (_ρ_ = 0.49, *p* = 0.05) in the CU-aCSF group, whereas at the Braak V (_ρ_ = 0.33, *p* = 0.05) in the MCI group.

### gBOLD activity links NbM degeneration to regional tau accumulation

The NbM-gBOLD correlation map in the CU-aCSF group was inversely similar to its gBOLD-tau correlation map (_ρ_ = –0.55, *p* = 1.15 × 10□□), with both showing the strongest correlations at early Braak stage regions, including the entorhinal cortex, parahippocampal gyrus, fusiform gyrus, and inferior and middle temporal cortices (**Fig. 3A**). Within the meta-temporal ROI, lower baseline gBOLD activation was associated marginally with greater tau accumulation at follow-up (_ρ_ = –0.47, *p* = 0.06; **Fig. 3B**), but the reverse relationship (i.e., baseline tau vs. longitudinal gBOLD change) was completely absent (_ρ_ = 0.03, *p* = 0.9; **Fig. S4A**). Smaller baseline NbM volume also predicated greater tau accumulation in this region (_ρ_ = –0.53, *p* = 0.03), but this association was no longer significant after controlling for gBOLD activations in these regions (_ρ_ = –0.27, *p* = 0.30; **Fig. 3C**). Similar patterns were observed in the MCI group, with the primary associations shifted towards advanced Braak regions. The NbM–gBOLD and gBOLD–tau correlation maps show inversely similar patterns (_ρ_ = –0.34, *p* = 0.005), with the strongest correlations found in the anterior and medial prefrontal cortex regions (**Fig. 3D**). Weaker baseline gBOLD activity in the Braak V ROI predicted greater longitudinal increases in tau accumulation (_ρ_ = –0.46, *p* = 0.007; **Fig. 3E**), while the reverse relationship was non-significant (_ρ_ = 0.16, *p* = 0.35; **Fig. S4B**). Similarly, smaller baseline NbM volume predicted greater tau accumulation in the Braak V ROI for the MCI group (_ρ_ = –0.38, *p* = 0.03), and this association became non-significant with adjusted for gBOLD activation (_ρ_ *= –0.26, p = 0.14;* **Fig. 3F**). Thus, baseline NbM volume and gBOLD activity are associated with baseline tau levels and its longitudinal changes in a spatially corresponding way, progressing from early Braak regions at the preclinical stage to more advanced Braak regions at the prodromal stage (**Fig. 3G**).

**Figure 3.**
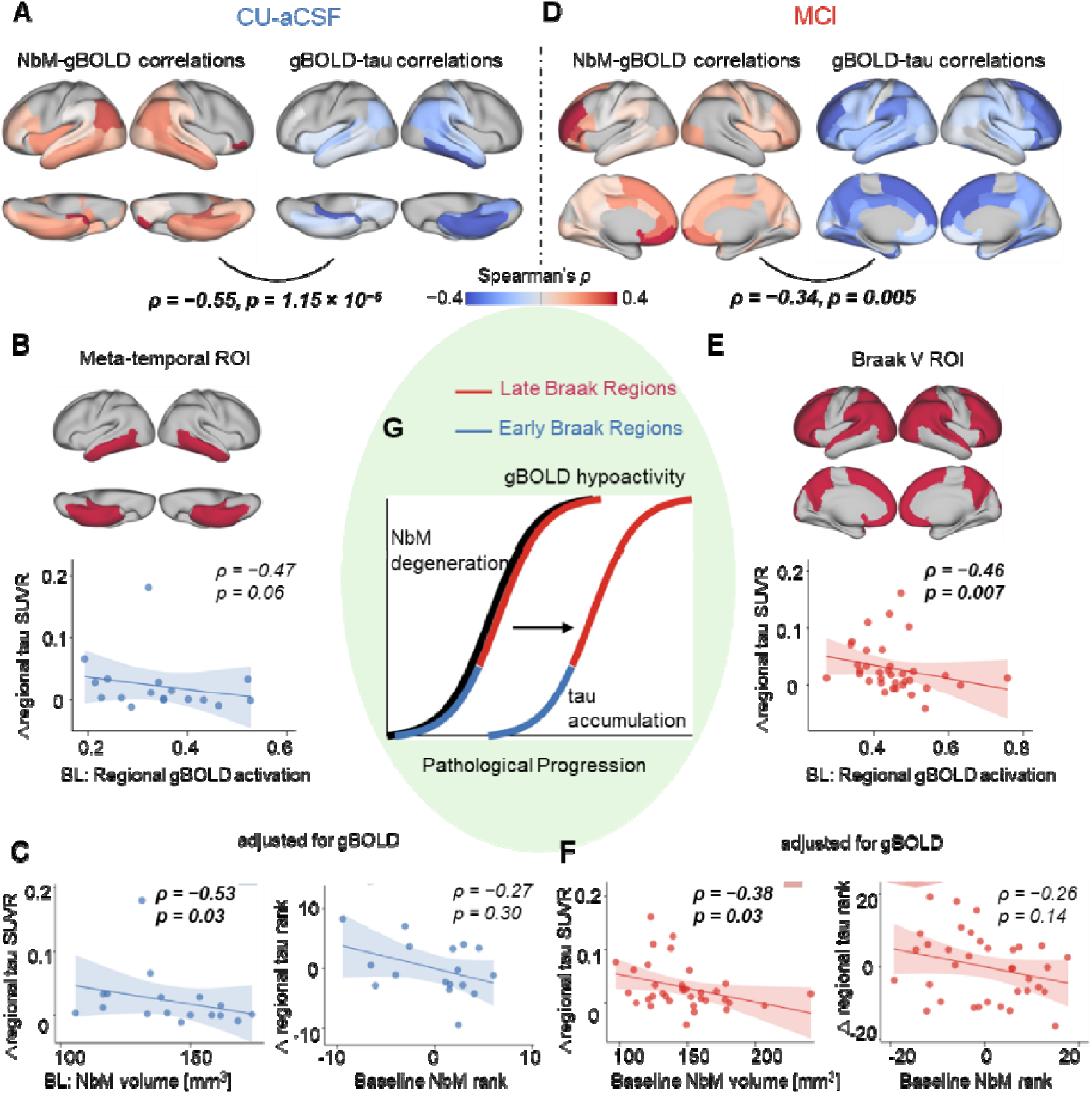
Three-way relationships between NbM volume, gBOLD activity, and cortical tau accumulation. (**A**) In the CU-aCSF group, the NbM-gBOLD correlation map (left) is inversely correlated with the gBOLD-tau correlation map (right) (ρ = –0.55, p = 1.15 × 10□□), with thei strongest positive (left) and negative (right) correlations overlapped at the early Braak regions. *(B)* Weaker baseline gBOLD activation in the meta-temporal ROI was marginally associated with more subsequent tau accumulation locally at the same regions (ρ = –0.47, p = 0.06). (**C**) Smaller baseline NbM volume predicted greater tau accumulation in the meta-temporal ROI (ρ = –0.53, p = 0.03), but the association was no longer significant after controlling for regional gBOLD activation (ρ = –0.27, p = 0.30). (**D**) In the MCI group, the spatial similarity between the NbM-gBOLD and gBOLD-tau correlation maps remained (ρ = –0.34, p = 0.005), but their most significant correlations progressed to more advanced Braak regions. (**E**) In the Braak V ROI, weaker baseline gBOLD activation predicted greater longitudinal tau accumulation locally (ρ = –0.46, p = 0.007). (**F**) Smaller baseline NbM volume also predicted the tau increases in thi region (ρ = –0.38, p = 0.03), but their association became non-significance after adjusting for regional gBOLD activation (ρ = –0.26, p = 0.14). (**G**) Schematic illustration of the proposed disease-stage model: gBOLD hypoactivity links NbM degeneration to tau accumulation. The black curve: NbM degeneration; the left blue-red curve: gBOLD hypoactivity; the right blue-red curve: tau accumulation.

### gBOLD activity and tau pathology are related to cognitive performance

We next examined how gBOLD activity and tau pathology are related to cognitive measures, i.e., the Mini-Mental State Examination (MMSE). Stronger mean gBOLD activations were associated with higher MMSE scores in both CU-aCSF (_ρ_ = 0.55, *p* = 0.02) and MCI (_ρ_ = 0.62, *p* = 7.76 × 10□□) groups, whereas the association between mean cortical tau SUVR and MMSE scores was only significant in the MCI group (_ρ_ = –0.61, *p* = 1.14 × 10□□) (**Fig. 4A**-**4B**). Stratifying these correlations by different Braak ROIs revealed distinct patterns. The strongest gBOLD-MMSE association progressed from the Braak III-IV region in the CU-aCSF group to the Braak V region in the MCI group (**Fig. 4C**-**4D**). The tau-MMSE correlations also exhibited a progressing pattern from the meta-temporal region (only marginally significant, *p* = 0.06) in the CU-aCSF to the Braak III-IV regions in the MCI group (**Fig. 4E**-**4F**). Notably, for both groups, the peak gBOLD-MMSE associations appeared in more advanced Braak regions than the peak tau-MMSE associations (**Fig. 4G**-**4H**). No significant gBOLD–MMSE or tau–MMSE associations were observed in the CU-nCSF group (**Fig. S5**).

**Figure 4.**
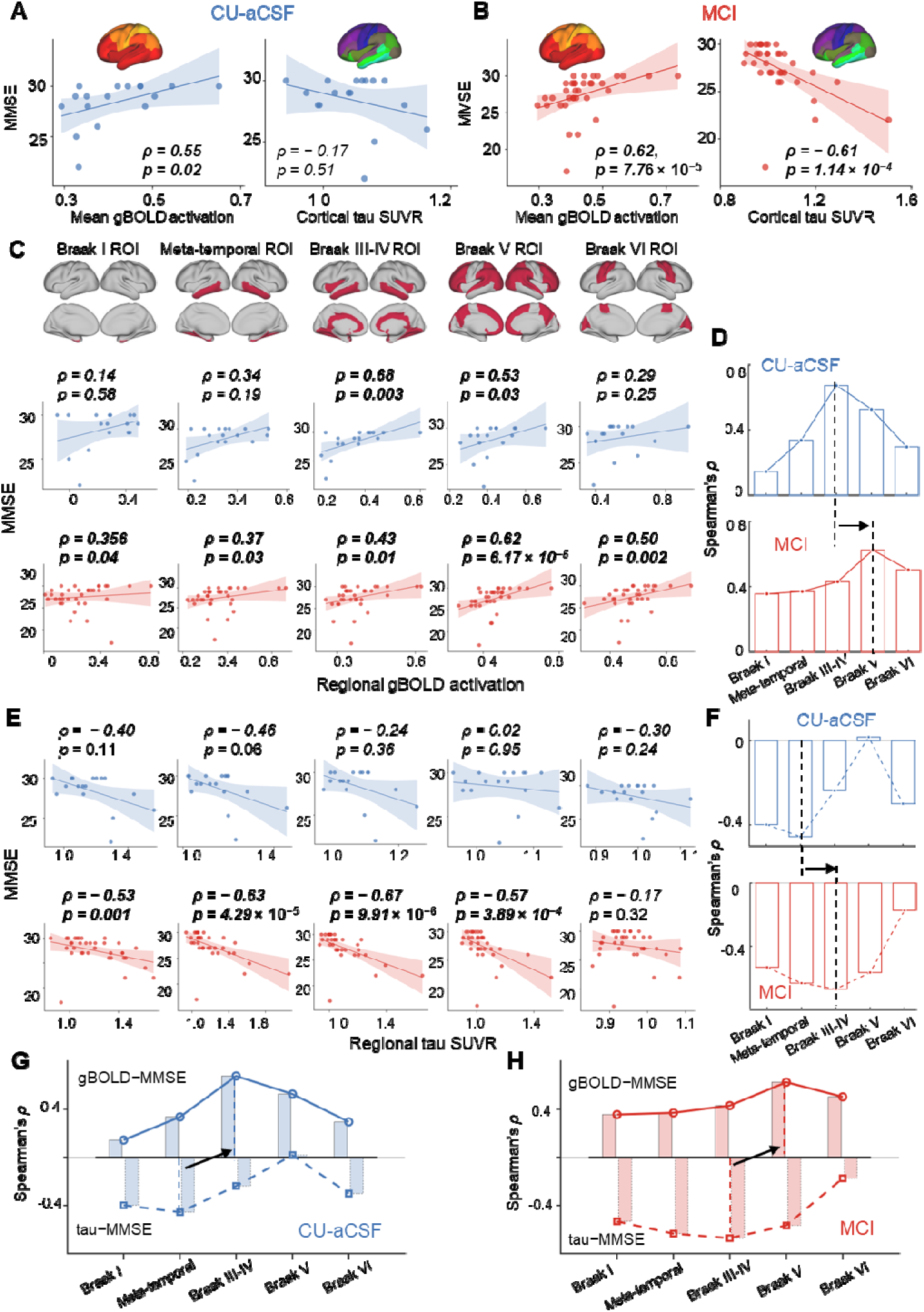
gBOLD activity and tau pathology are associated with cognitive measures. (**A**) higher MMSE scores were significantly correlated with weaker mean gBOLD activation (**A**; ρ = 0.55, p = 0.02) but not mean cortical tau SUVR (ρ = –0.17, p = 0.51) in the CU-aCSF group. (**B**) MMSE scores were associated with both mean cortical tau SUVR (ρ = –0.61, p = 1.1 × 10□□) and mean gBOLD activation (ρ = 0.62, p = 7.8 × 10□□) in the MCI group. (**C**) Regional analyses show that gBOLD–MMSE correlations peaked in the Braak III–IV regions for the CU-aCSF group (ρ = 0.68, p = 0.003) and in the Braak V regions for the MCI group (ρ = 0.62, p = 6.17 × 10^-5^). (**E**) Tau–MMSE correlations are the strongest in the meta-temporal regions for the CU-aCSF group (ρ = –0.46, p = 0.06) whereas in Braak III–IV regions for the MCI group (ρ = −0.67, p = 9.91 × 10^−6^). Bar plots summarize gBOLD-MMSE (**D**) and tau-MMSE (**F**) associations across Braak regions for each group. Region-specific comparisons of gBOLD-MMSE and tau-MMSE correlations in the CU-aCSF (**G**) and MCI (**H**) groups.

## Discussion

Using multimodal data from the ADNI3 project, we identified systematic relationships among NbM degeneration, disrupted infra-slow gBOLD activity, cortical tau accumulation, and cognitive performance in early AD. We found that smaller NbM volume was associated with spatially co-localized reductions in gBOLD activity and increases in tau accumulation. These NbM-related changes progressed from early Braak regions in preclinical CU-aCSF subjects to advanced Braak areas in prodromal MCI subjects. Predictive modeling of longitudinal data further supports a directional model in which gBOLD hypoactivity associated with smaller NbM predicts subsequent increases in cortical tau pathology. Both gBOLD reduction and tau deposition were also associated with lower cognitive scores in these at-risk participants, but cognition-related gBOLD changes appeared in more advanced Braak regions than tau-related gBOLD changes for the both CU-aCSF and MCI groups. Together, these findings suggest a potential role of infra-slow gBOLD activity as a functional mediator that amplifies the effect of subcortical cholinergic degeneration on cortical tau pathology progressively across Braak stages from the early preclinical to prodromal stages of AD.

The connection between the cholinergic system and the formation of cortical plaques and neurofibrillary tangles was recognized long before the amyloid-β cascade hypothesis. The cholinergic pathogenesis of neuritic plaques is not only supported by the presence of acetylcholinesterase activity within plaques^39^ but also by the topological correspondence between cortical plaque deposition and NbM neuronal loss^16^. Subsequent work further showed that the loss of cortical cholinergic fibers aligns more strongly with neurofibrillary tangle burden than with Aβ deposits^18^. Importantly, the tau pathology appears in the NbM at very early preclinical stages^19–22^. Consistent with these histological results, *in vivo* neuroimaging studies have repeatedly reported the association between NbM degeneration and tau pathology at preclinical AD stages^23,40–43^. The NbM-tau association we observed is consistent with these early observations, confirms this relationship in the early preclinical AD stage, and further reveals its topological progression during the prodromal stage. While prior work reported baseline tau predicted NbM degeneration^42^, we observed only weak evidence for this temporal relationship in the CU-aCSF group. Instead, baseline NbM volume consistently predicted subsequent tau accumulation in both the CU-aCSF and MCI groups, though in different sets of brain regions. The finding aligns with recent evidence that NbM degeneration precedes the cortical spread of Alzheimer’s pathology/degeneration, even in the transentorhinal regions^24,25^ previously thought to be the initial site of AD pathology^44,45^, challenging the traditional view that the NbM changes as a downstream or secondary consequence of cortical pathology^15,46,47^. The early presence of tau pathology in the NbM, coupled with its known widespread cortical projections^48,49^, suggest it may act as a cholinergic “amplifier”, if not an “initiator”, that propagates Aβ/tau pathology to the cortex^50^. However, the mechanism by which the subcortical NbM pathology propagates to cortical regions remains to be elucidated.

Recently identified infra-slow global brain activity may represent such a missing link between the NbM degeneration and cortical Aβ/tau pathology. This global brain activity is manifested as highly structured gBOLD waves propagating across cortical hierarchies in human fMRI and monkey electrocorticography (ECoG)^26–28^, whereas as stereotypic cascades of spiking sequences in large-scale neuronal recordings ^29^. Converging evidence indicated a tight link between gBOLD activity and the cholinergic arousal system: this activity is coupled to arousal modulations reflected in pupil size and slow wave activity and is also characterized by strong and specific de-activation in the NbM^30,51,52^. Importantly, pharmaceutical de-activation of the NbM in one hemisphere effectively suppressed gBOLD activity in the ipsilateral cortex in monkeys^31^. Recent PET–MRI evidence in humans directly associated gBOLD activity with cortical cholinergic density^37^. Interestingly, age-related declines in gBOLD activity and NbM volume appeared to follow similar trajectories, with reductions beginning around age 55^53,54^. Our findings substantiate the gBOLD-cholinergic relationship and extend prior observations by showing the direct associations between gBOLD activity and NbM volume.

Meanwhile, converging evidence also linked gBOLD activity to cortical amyloid/tau accumulation, an association often interpreted from the perspective of brain clearance. The potential role of gBOLD activity in brain waste clearance was initially suspected due to its sleep-dependent coupling with CSF inflow^55^, a process central to a newly proposed glymphatic clearance pathway^56–58^. Subsequent studies found significant associations between the gBOLD-CSF coupling strength with both Aβ and tau pathology^32,34,59^. Importantly, topological changes in gBOLD wave dynamics were further found to explain the spreading pattern of Aβ in the cortex^33^. The role of gBOLD activity in cerebral clearance is likely mediated by the coordinated actions of subcortical neuromodulatory nuclei, particularly the NbM and the brainstem locus coeruleus (LC)^36^, on cerebral arteries to promote infra-slow vasomotions^35^. This hypothesis is supported by recent animal research showing that cholinergic lesions lead to reduced gBOLD-CSF coupling, arterial pulsations, and eventually impaired clearance function^37^.

Our results expanded these early findings by linking gBOLD and tau changes to NbM degenerations and showing their associated progression across Braak-staged cortical regions from preclinical to prodromal stages of AD. The spatial pattern of gBOLD activity, i.e., the gradually increased co-activation levels across Braak regions, already hinted its potential connection to tau spreading and Braak stages. This negative spatial correlation with the tau map mirrors the classic observation that regions with sparse cholinergic innervation exhibit dense Aβ plaques deposition^14,15^, a pattern cited to support for a secondary role of the NbM in AD pathology^60^. However, such inverse relationship aligns well with a clearance-based interpretation, as sparse cholinergic inputs could make a region more susceptible to clearance impairment. Cross-subject gBOLD-tau correlations provide stronger evidence for their relationship and further demonstrate that the progression of this association from early Braak regions in the preclinical stage to more advanced Braak areas in the prodromal stage. Interestingly, longitudinal predictive modeling suggested a unidirectional relationship, in which baseline gBOLD activity consistently predicted later cortical tau accumulations in both preclinical and prodromal cohorts and contributed to the association between baseline NbM volume and subsequent tau increases. The finding thus supports a possible role of gBOLD activity in mediating the effects of NbM degenerations on cortical tau pathology.

Both gBOLD activity and cortical tau are correlated with cognitive performance, even among cognitively unimpaired subjects (CU-aCSF). The gBOLD correlation with cognition may reflect its close relationship to the cholinergic system, which plays critical roles in attention regulation and synaptic plasticity^61–63^, and thus cognitive and memory functions. In fact, the infra-slow global brain activity, i.e., spiking cascades in mice and gBOLD activity in humans, has been linked to memory processes. These include hippocampal ripples and replays in mice^29,64^ and memory retrieval and cognitive measures in humans^51,65,66^. In clinical cohorts, the gBOLD-CSF coupling has also been associated with cognitive decline in both AD/MCI and PD patients^59,67^. An interesting observation of the present study is that, in both the CU-aCSF and MCI groups, the cognition score is correlated with gBOLD activity in more advanced Braak regions than with tau pathology, even in brain areas with non-significant NbM-gBOLD correlation, such as Braak III-V for the CU-aCSF group. One possible explanation is that NbM neurons projecting to these regions are already functionally impaired, disrupting gBOLD and cognition but not yet causing measurable NbM atrophy or local tau accumulation.

Our NbM-related findings do not rule out the contributions of other neuromodulatory nuclei, particularly the brainstem LC, to gBOLD and tau changes during AD progression. In fact, gBOLD waves are coupled to spontaneous pupil dilations and also accompanied by LC de-activations at early phases^26^. Meanwhile, both anatomical connectivity and the pattern of tau accumulation suggest that the LC may act as an even more upstream node than the NbM in AD progression^68–70^. Nevertheless, quantifying LC volume is extremely challenging, if not impossible, with current ADNI imaging data. Future work using specialized neuromelanin-sensitive MRI or high-resolution brainstem imaging will be essential to quantify LC volume/contrast and clarify its role in gBOLD activity and cortical tau pathology.

In summary, we found that NbM degeneration is associated with reduced gBOLD activity and increased tau accumulation in early Braak regions during the preclinical AD stage, and these changes progress to more advanced Braak areas in the prodromal stage. These findings support the potential role of infra-slow gBOLD activity as a neural mediator that amplifies early subcortical NbM dysfunction to cortical tau pathology.

## Data Availability

All data used in this study are publicly available through the ADNI website (https://adni.loni.usc.edu/), including subjects characteristics, Aβ42 and p-tau in CSF, rsfMRI, structural MRI, and Mini-Mental State Examination (MMSE) scores were obtained from the same study visit defined by ADNI. The file “UC Berkeley—AV1451 analysis [ADNI1,GO,2,3] (version: 2022-04-26)” was used to provide the tau-PET SUVR. CSF Aβ42 and CSF p-tau data for our cohort were obtained from “UPENN CSF Biomarkers Roche Elecsys [ADNI1,GO,2,3]”. The Mini-Mental State Examination (MMSE) scores were obtained from “Mini-Mental State Examination (MMSE) [ADNI1,GO,2,3,4]”. The DKT-68 atlas was obtained from the website https://surfer.nmr.mgh.harvard.edu/fswiki/CorticalParcellation.

## Code Availability

Data and code used to generate the main results are available upon request.

## Acknowledgments

This work was supported by the Brain Initiative award (1RF1MH123247-01) and the NIH R01 award (1R01NS113889-01A1). Data collection and sharing for this project was funded by the Alzheimer’s Disease Neuroimaging Initiative (ADNI) (National Institutes of Health Grant U01 AG024904) and DOD ADNI (Department of Defense award number W81XWH-12-2-0012). ADNI is funded by the National Institute on Aging, the National Institute of Biomedical Imaging and Bioengineering, and through generous contributions from the following: AbbVie, Alzheimer’s Association; Alzheimer’s Drug Discovery Foundation; Araclon Biotech; BioClinica, Inc.; Biogen; Bristol-Myers Squibb Company; CereSpir, Inc.; Cogstate; Eisai Inc.; Elan Pharmaceuticals, Inc.; Eli Lilly and Company; EuroImmun; F. Hoffmann-La Roche Ltd and its affiliated company Genentech, Inc.; Fujirebio; GE Healthcare; IXICO Ltd.; Janssen Alzheimer Immunotherapy Research & Development, LLC.; Johnson & Johnson Pharmaceutical Research & Development LLC.; Lumosity; Lundbeck; Merck & Co., Inc.; Meso Scale Diagnostics, LLC.; NeuroRx Research; Neurotrack Technologies; Novartis Pharmaceuticals Corporation; Pfizer Inc.; Piramal Imaging; Servier; Takeda Pharmaceutical Company; and Transition Therapeutics. The Canadian Institutes of Health Research is providing funds to support ADNI clinical sites in Canada. Private sector contributions are facilitated by the Foundation for the National Institutes of Health (www.fnih.org). The grantee organization is the Northern California Institute for Research and Education, and the study is coordinated by the Alzheimer’s Therapeutic Research Institute at the University of Southern California. ADNI data are disseminated by the Laboratory for Neuro Imaging at the University of Southern California.

The authors of this work recognize the Penn State Institute for Computational and Data Sciences (RRID:SCR_025154) for providing access to computational research infrastructure within the Roar Core Facility (RRID: SCR_026424).

## Competing interests

The authors declare no competing interests.

## Notes

### Competing Interest Statement

The authors have declared no competing interest.

